# Influence of nutritional tyrosine on cognition and functional connectivity in healthy old humans

**DOI:** 10.1101/450650

**Authors:** Christian Hensel, Maxi Becker, Sandra Düzel, Ilja Demuth, Kristina Norman, Elisabeth Steinhagen-Thiessen, Jürgen Gallinat, Ulman Lindenberger, Simone Kühn

**Affiliations:** University Clinic Hamburg-Eppendorf, Clinic and Policlinic for Psychiatry and Psychotherapy, Martinistraße 52, 20246 Hamburg, Germany; Max Planck Institute for Human Development, Center for Lifespan Psychology, Berlin, Germany; Charité – Universitätsmedizin Berlin, corporate member of FreieUniversität Berlin, Humboldt-Universitätzu Berlin, and Berlin Institute of Health, Department of Endocrinology, Diabetes and Nutrition (including Lipid Metabolism), Germany; Berlin-Brandenburg Center for Regenerative Medicine (BCRT), Charité University Medicine Berlin, Germany; Charité – Universitätsmedizin Berlin, corporate member of FreieUniversität Berlin, Humboldt-Universitätzu Berlin, and Berlin Institute of Health, Research Group Geriatrics, Germany; German Institute of Human Nutrition Potsdam - Rehbrücke, Department of Nutrition and Gerontology, Nuthetal, Germany; Max Planck Centre for Computational Psychiatry and Ageing Research, London, United Kingdom

**Keywords:** Tyrosine, working memory, resting state, functional connectivity

## Abstract

Tyrosine is precursor for monoamine neurotransmitters such as dopamine (DA), which is one of the key neurotransmitters in the frontostriatal network and of crucial relevance for mental disorders. Recent research reported that high dose tyrosine application resulted in increased brain DA synthesis, which is consistent with the observation of positive associations between daily tyrosine intake and cognitive test performance. In the present study, we investigated the associations between working memory (WM) dependent tasks and self-reported nutritional tyrosine intake within a large group of healthy elderly humans (286 subjects) by additionally including brain functional data. We observed a negative correlation between tyrosine intake and resting-state functional connectivity (rsFC) between the striatum (putamen) and the prefrontal cortex. That is to say, we found higher rsFC in individuals consuming less tyrosine per day. At the same time, this increased rsFC or hyperconnectivity was associated with lower WM performance. These findings suggest that lower or insufficient supply of tyrosine might result in dysfunctional connectivity between striatal and frontal regions leading to lower WM capacity in healthy elderly humans.

## Introduction

In times of superannuation of the population, it becomes ever more important for people to stay healthy over the course of their whole lifetime. One main aspect of age-related health problems is cognitive deterioration. While most people think it is possible to prevent cognitive impairment due to ageing, a lot of them are not sure how to do so (Vaportzis & Gow, 2018). Alongside with physical exercise and improvement in healthcare, nutrition has gotten into focus as a contributor in achieving this goal. Over the past few years, we gained a better insight of several different nutrients and their role in the healthy human brain. One of them is tyrosine, an essential amino acid, which is present in protein rich food (e.g. almonds, eggs, salmon). It serves as the precursor of catecholamine neurotransmitters (CA) (dopamine, epinephrine, norepinephrine). The rate-limiting step in the synthesis of CA is the enzyme tyrosine-hydroxylase (TH). Besides various other factors, research suggests an influence of precursor availability on the activity of TH in mammalian brains (Wurtman et al. 1974) due to TH not being fully saturated most of the time (Carlsson & Lindqvist, 1978). Substrate availability corresponds to tyrosine levels in the brain. Brain levels of tyrosine may vary as a function of its availability in the blood (Fernstrom & Fernstrom, 2007),due to a direct link between blood and brain concentrations of tyrosine. For this reason, nutrition might have an impact on brain CA synthesis and activity, due to nutritional intake determining blood levels of tyrosine (Moja et al., 1996).

Synthesis and metabolism of CA neurotransmitters, in particular dopamine (DA), has a variety of effects on the central nervous system. Evidence provided by pharmacological intervention shows its importance in cognitive processes. Early animal studies have shown that depletion of DA in regions of the prefrontal cortex (PFC) had effects on working memory (WM) in monkeys similar to surgical ablation of the same area (Brozoski et al., 1979). Pharmacological intervention using levodopa (L-Dopa, a DA precursor) reversed those drug-mediated effects of dopamine depletion. Subsequent animal studies showed similar effects of DA modulation in the PFC and impaired cognitive function (Arnsten, 1989; Watanabe et al., 1997). Research in healthy human volunteers provided further evidence for the relationship between DA and cognition. Administration of DA D2 receptor antagonists to reduce DA transmission led to impaired spatial working memory performance (Mehta et al., 1999), while low doses of an agonist improved performance (Luciana et al., 1998). However, this correlation is not linear, meaning higher doses of a DA agonist or DA brain levels itself do not always result in better performance. In fact, the relationship resembles more an inverted U-shape, meaning too low or too high levels can impair cognitive control and effects are highly dependent on the basal supply of neurotransmitters (Cools &D’Esposito, 2011). Therefore, maintaining an optimal level of DA seems to be crucial. Besides direct manipulation of transmitter signaling in drugs, manipulation of the precursor tyrosine had similar effects on cognition. For example, depletion of tyrosine led to a lower release of DA and impaired cognitive performance in healthy volunteers (Harmer et al., 2001). This shows the importance of a constant supply of tyrosine for maintaining DA synthesis and signaling. In addition, former research has shown tyrosine intake can also enhance WM performance and reverse negative effects of stress in WM tasks (Banderet & Lieberman, 1989; Shurtleff et al. 1994; Thomas et al., 1999; Deijen et al., 1999). This suggests tyrosine to be especially effective in times of high demand or lower basal levels of DA (e.g. under stressful conditions). Furthermore, ingestion of tyrosine rich food improved WM capacity and task-related functional connectivity between the PFC and striatum compared to food missing tyrosine (Nagano-Saito et al., 2008). These findings show that tyrosine-mediated effects are not limited to altering cognition but also affect functional brain connectivity. Another example for decline of the DAergic system is a pathological condition known as Parkinson’s disease (PD). Here dopamine depletion within the striatum leads to loss of cognitive and motor abilities. Impairments linked to this condition are rather specific for frontostriatal cognitive function (Lange et al., 1992; Owen et al., 1992; Lees & Smith, 1983). Additionally, as shown by neuroimaging studies, lower activity within the PFC and striatum can be observed in PD patients, while performing a WM capacity depending cognitive task (Owen et al., 1998; Lewis et al., 2003). At the same time, studies on healthy humans have provided evidence for a correlation between higher DA synthesis in the striatum and higher WM capacity (Cools & D’Esposito, 2011). That is to say, WM performance may serve as a predictor for striatal DA synthesis capacity (Cools et al., 2008). Interestingly research has also shown an effect of nutritional tyrosine on DA release in striatal areas (Montgomery et al., 2003). That is why tyrosine blood levels might influence the DAergic system by changes of DA in prefrontal and striatal regions, due to precursor availability, leading to changes in WM task performance and functional connectivity within the frontostriatal network. So far, most research focused on administration of relatively high doses of tyrosine immediately before the tests. To investigate real-life effects of differences in daily dietary tyrosine intake, we recently monitored nutritional behavior as well as cognitive function of healthy adults. The results showed associations between higher everyday tyrosine intake and better WM capacity in younger and older adults (Kühn et al., 2017).

Another rather natural cause of changes in the DAergic system accompanied by cognitive decline is age. Evidence shows an age-related reduction in DA activity in the brain, associated with the decline of motor and cognitive functions (Volkow et al. 1998). Precisely, striatal impairment in DA metabolism and signaling correlates with cognitive deterioration in healthy elderly humans (Vernaleken et al., 2007; Erixon-Lindroth et al., 2005; Bäckman et al., 2000). In contrast, research suggests DA synthesis capacity to be upregulated with age, possibly as a tool to compensate for reduced signaling (Braskie et al., 2008). Thus, previously described effects of tyrosine availability on cognition and its effectiveness in times of high demand, might be especially prevalent in senescent adults, due to age-related decline of the DA metabolism and therefore higher DA demand.

Given the evidence stated above, we hypothesized a positive effect of a tyrosine rich diet on brain connectivity and WM capacity. To investigate this, we first analyzed data of nutritional behavior and functional brain connectivity. We assessed the everyday intake of tyrosine via a food frequency questionnaire. Subsequently, we analyzed resting-state functional connectivity (rsFC) in fMRI in association to tyrosine intake. Functional connections between PFC and striatum are in close relationship with working memory and can be monitored by rsFC (Di Martino et al., 2008). Previous work has shown that pharmacological manipulation of DA, as the one described above, influences rsFC of the frontostriatal network (Cole et al., 2013a) and other resting-state networks (Cole et al., 2013b; Haaker et al., 2016). A recent study revealed hyperconnectivity of striatal and prefrontal regions in patients suffering from early stages of PD. Administration of L-DOPA mitigated this abnormal rsFC(Kwak et al., 2010). Therefore, under the assumption of dietary tyrosine influencing the DAergic system, we expected to see a correlation between individual tyrosine intake and rsFC of the PFC and the striatum. We focused our research on a group of healthy elderly humans (older than 60 years) of the Berlin Aging Study II (https://www.base2.mpg.de/en), since they display higher variability of WM performance due to age-related cognitive decline. Furthermore, we expected effects of dietary tyrosine to be more prevalent in old age, due to higher demand of DA supply in age. Additionally, a broad battery of cognitive taskswas used to determine WM capacity and to search for associations between daily dietary tyrosine intake, brain connectivity and cognitive performance.

## Methods

### Participants

We recruited participants within the Berlin Aging Study II (BASE-II) (for additional information see Bertram et al. (2014) and Gerstrof et al.(2016)). Out of the original 2200 participants, fMRI data of 332 subjects older than 60 years was available. Six subjects were excluded due to low scan quality and morphological anomalies as well as another 40 subjects due to missing data in nutrition or cognitive assessment, respectively. This resulted in a final sample size of 286 participants. Subjects were between 60-81 years old (age in years, M=69.6, SD=3.9, 116 female). None of the participants had a history of head injuries, medical (e.g., heart attack), neurological (e.g., epilepsy), or psychiatric disorders (e.g., depression) or were on medication associated with these condition (e.g. antiepileptic, neuroleptic, antidepressant drugs), which may have affected memory function.

### Procedure

On average, nutritional data was acquired two years prior to cognitive assessment (mean difference in days = 719, *SD* = 435, range = 131 – 1960) and MRI-scans (mean difference in days = 824, *SD* = 430, range = 1 – 767). MRI-scans took place after the cognitive assessment, on average 3.5 months (mean difference in days = 106, *SD* = 124).

### Nutrition assessment

For the assessment of nutritional behavior, we used the European Prospective Investigation into Cancer and Nutrition (EPIC) food frequency questionnaire (FFQ) (Boeing et al., 1997). This questionnaire assesses the dietary intake of 148 food items over the course of the last 12 months. The portion size of each item was indicated by line drawings and the participants had to rate their frequency of consumption. Then, using values provided by the Federal Coding System, we determined the nutrients concentration for every food item of the FFQ. For the following analysis, we used the average amount of tyrosine and the average amount of food consumed per day in grams.

### Cognitive assessment

The cognitive battery of BASE-II included the assessment of multiple tasks. Here, we focused on the cognitive ability of WM indicated by Letter Updating, Spatial Updating, Serial Recall and Number-N-Back. For all tasks we used accuracy as our main indicator for performance.

#### Letter Updating Task (LU)

Subjects were presented with 7, 9, 11 or 13 letters in a sequence. After the end of a sequence, the subjects had to report the last 3 letters of the sequence in correct order by pressing buttons corresponding to A, B, C, and D.

#### Spatial Updating (SU)

Participants had to remember and update the position of a blue dot within 3×3 grids. First, two grids were displayed for 4000ms. Within each of those grids a blue dot was presented in one of the 9 locations. The participants had to remember and update the location according to shifting operations, which were indicated by arrows appearing below the corresponding grids. Arrows were displayed for 2500ms. After six operations, the grids reappeared and the subjects had to click on the resulting end position. The same task was repeated with three instead of two grids shown simultaneously.

#### Serial Recall (SR)

The participants were instructed to memorize three lists consisting of twelve words each, which were presented via headphones. Simultaneously, the serial position of each word was presented via numbers on a computer screen. After list presentation, the subjects had to recall the items in forward (1 to 12) or backward (12 to 1) order.

#### Number-N-Back Task (NNB)

Participants had to decide whether the current stimulus matches the stimulus shown three steps earlier in the sequence. The stimuli were one-digit numbers (1–9) presented consecutively in 3 visible squares. Subjects made their decision via button-box presses (left/right index finger on white and black button).

#### Working Memory Latent factor

In order to find a more robust measure for general working memory capacity(to test for relations with tyrosine consumption and differences in functional brain connectivity),we computeda latent factor for WM. The latent factor was determined by the shared covariance of all four above mentioned WM tasks (SU, LU, SR, NNB) ina confirmatory factor analysis (CFA). The CFA was carried out using R (RStudio Team, 2016) and the lavaan package (Yves Rosseel, 2012). Due to chi-square being very sensitive to sample size,a goodness of model fit is commonly indicated by additional fit indices like Root Mean Square Error of Approximation (RMSEA), Standardized Root Mean Square Residual (SRMR) and Comparative Fit Index (CFI). Accepted thresholds indicating good model fit are RMSEA<= .05, SRMR<= .1 and CFI >= .95(Schermelleh-Engel et al. 2014; Hu and Bentler 1999).

### Scanning Procedure

Brain images were collected with a Siemens Tim Trio 3T scanner (Erlangen, Germany) using a 12-channel head coil. A T1-weighted magnetization prepared gradient-echo sequence (MPRAGE) based on the ADNI protocol (TR=2500ms; TE=4.77ms; TI=1100ms, acquisition matrix=256×256×176; flip angle = 7˚; 1×1×1mm^3^voxel size) was used to obtain structural images. Whole brain functional resting state images were collected over a period of 10 minutes by using a T2*-weighted EPI sequence sensitive to BOLD contrast (TR=2000ms, TE=30ms, image matrix=64×64, FOV=216mm, flip angle=80°, slice thickness=3.0mm, distance factor=20%, voxel size 3×3×3mm^3^, 36 axial slices. Participants were instructed to fixate a fixation cross and relax during data acquisition.

### Functional connectivity analysis using resting-state data

The following steps of preprocessing functional resting-state images and statistical analysis (with time series between ROIs) were performed using CONN-fMRI functional connectivity toolbox, version 16.b (www.nitrc.org/projects/conn) (Whitfield-Gabrieli& Nieto-Castanon 2012) based on SPM12 (www.fil.ion.ucl.ac.uk/spm/). In order to reduce the fluctuation of MRI signal in the initial stages of the scan, the first 10 functional images of every subject were discarded, resulting in 290 images per subject.

#### Preprocessing

At first, we performed spatial preprocessing of functional and structural images. This included slice-timing correction, motion realignment, coregistration, gray and white matter (WhM) as well as cerebrospinal fluid (CSF) structural segmentation, normalization of functional and structural scans to Montreal Neurological Institute (MNI) space and smoothing with an 8-mm FWHM-Gaussian filter, using the SPM12 default parameter choices.

The subjects’ estimated motion parameters, the global BOLD signal and BOLD signals in WhM and CSF were used as covariates to attenuate the impact of motion and physiological noise. We used the implemented CompCor approach (Behzadi et al., 2007) including regression of six realignment parameters as well as their first temporal derivatives and regression of WhM and CSF. After a linear detrending, the resulting time series were additionally band-pass filtered between 0.1 and 0.01 Hz to remove low-frequency drifts and physiological high-frequency noise.

#### First-level analysis

Using the standard atlas provided by the CONN toolbox (containing 132 ROIs, 106 cortical and subcortical areas from the FSL Harvard-Oxford Atlas and 26 cerebellar areas from the AAL Atlas) we computed functional connectivity between each pair of ROIs (ROI-to-ROI analysis).

#### Second-level analysis

To investigate the effects of dietary tyrosine, we used the average intake in grams per day (as stated above) and checked for differences in rsFC depending on the amount of tyrosine. We executed a whole brain analyses by correlating the functional connectivity of every ROI pair with the respective tyrosine value while correcting for multiple comparison (p<= .05, FDR corrected). Age, sex, years of education and average amount of food in grams per day were used as covariates of no interest.

#### Functional connectivity values

For an additional investigation of correlations between rsFC (between our ROIs stated above) and WM performance, we extracted the connectivity values between each pair of ROIs for every subject using the CONN output generated during the first-level analysis.

### Behavioral data analysis

The obtained data was processed using Statistical Package for Social Science (SPSS, IBM Analytics) version 21. To test our hypothesis of dietary tyrosine intake predicting WM, we carried out multiple linear regression analysis, using average daily tyrosine intake and single subject connectivity values (as mentioned above) as predictors for WM. We used age, sex, years of education and average food intake per day as additional predictors (covariates of no interest). The threshold for all correlation magnitudes was *p* = 0.05.

Furthermore, we tested whether the effect of daily tyrosine intake on WM is mediated in the brain by differences in functional connectivity between striatum and prefrontal cortex. Therefore, we conducted a mediation analysis, as implemented in SPSS macro PROCESS (Hayes, 2013) to test for direct and indirect effects of tyrosine on WM mediated by rsFC. Significance of the indirect effect is indicated by the 95% confidence interval (CI) derived from 5000 bootstrap resamples. Mediation analysis was also controlled for age, sex, years of education and average food intake per day.

## Results

The participant group consumed an average of 2.84 g of tyrosine (*SD* = 0.93, range = 1.2 –6.5 g) and an average of 81.89g of protein (*SD* = 26.8, range = 36 – 200 g) per day. Their average daily food intake was 1389 g (*SD* = 405, range = 573 – 2745 g). We observed a significant negative correlation between the subjects’ age and their performance in the LU task [*r*(282) = -.117, p = .05], SU task [*r*(282) = -.184, p = .002] and NNB task [*r*(282) = -.132, *p* = .027]. The SR task did not show any significant association with age. In contrast, we saw a significant positive correlation between daily tyrosine intake and performance in the SU task [*r*(280) = .169, *p* = .004] and the SR task [*r*(280) = .188, p = .002], but not in the NNB and LU task (*see Table 1*). Meanwhile we did not observe any significant age related differences in daily tyrosine consumption (*p*> .3) and overall food intake (*p*> .7) within our group.

**Table 1:** Mean task-performance and correlation to age and tyrosine intake

| Cognitive Task | Participants' mean accuracy (%)<sup>a</sup> (*SD*) | Correlation with age (*r*-value)<sup>b</sup> | Correlation with tyrosine (*r*-value)<sup>c</sup> |
| --- | --- | --- | --- |
| Letter updating (LU) | 80.7 (22.2) | -.117** | .054 |
| Spatial updating (SU) | 51.3 (21.8) | -.184** | .169** |
| Serial recall (SR) | 35.1 (15.7) | -.030 | .188** |
| Number N-back (NNB) | 69.4 (17.7) | -.132** | .110* |

<sup>a</sup>participant's accuracy/correct answers in percent (of maximum possible)
<sup>b</sup>controlled for sex and years of education
<sup>c</sup> controlled for sex, age, overall food intake and years of education
\* $p < .1$ ; \*\* $p < .05$

### Latent factor for Working Memory

To define our latent factor capturing WM ability, we used test results for NNB, SR, SU, LU as stated above. In the resulting model, all variables loaded in the same direction (positive). Standardized factor loadings were within moderate range indicating that all tasks contributed significantly to the latent factor (NNB = 0.77, SU = 0.70, SR = 0.63, LU = 0.49). The exact test statistic did not reach significance [χ^2^ = 0.985, *p* = 0.611], but the practical fit indices [CFI = 1.0, RSME = < .001 and SMRE = .009] suggested a good fit of the model to our data. (*see Figure 1*).

**Figure 1:**
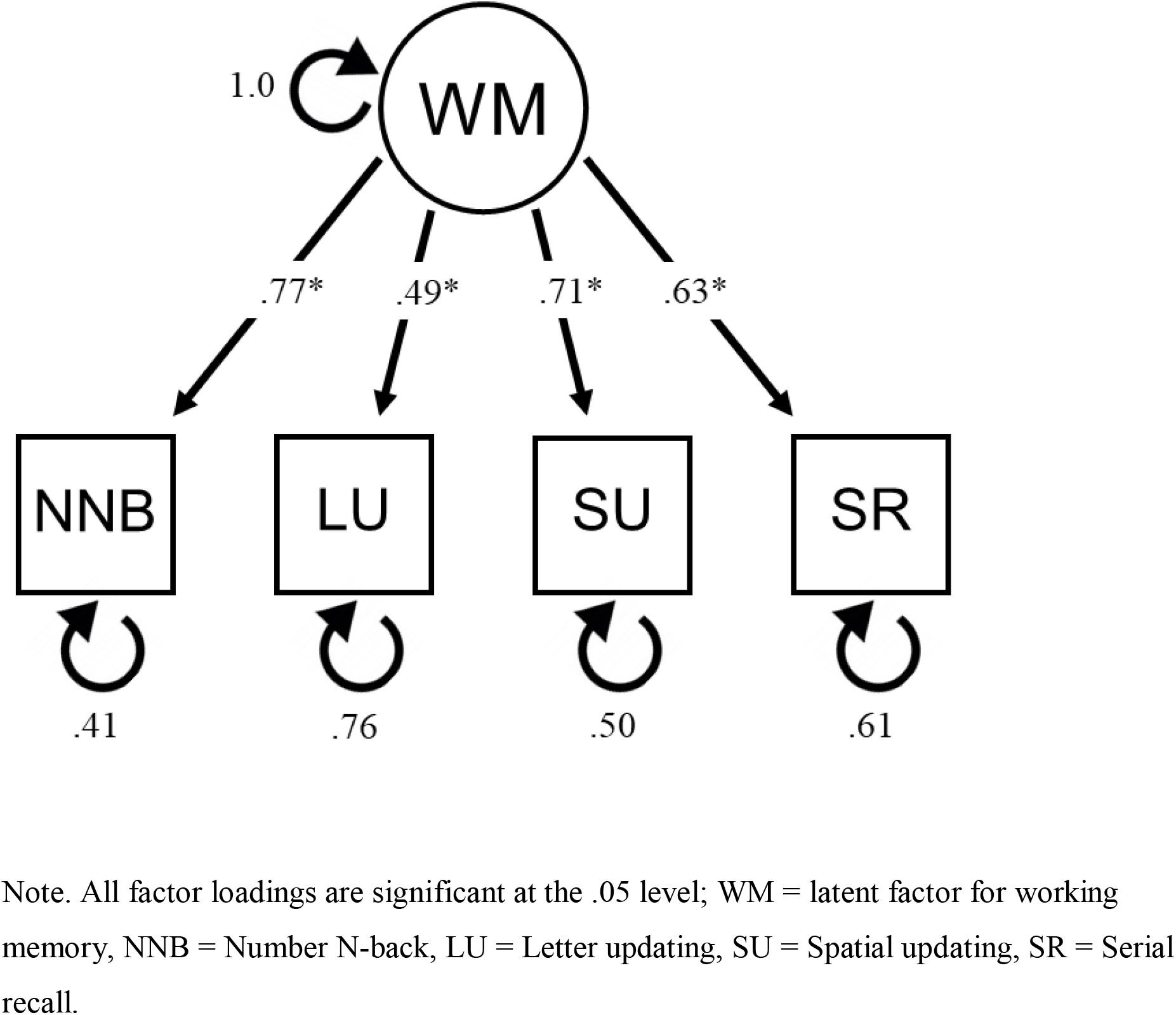
Measurement model for the latent factor working memory (WM)

### Functional connectivity analysis

Although we had a specific hypothesis focusing on prefrontal and striatal areas of the brain, we first performed a whole-brain ROI to ROI analysis to test for the robustness of the results. This whole-brain analysis revealed a significant negative correlation of daily tyrosine intake in gram and rsFC between the right putamen and left IFG pars opercularis [*β* = -0.40, *t*(280) =-3.63, *p-FDR* = .044] controlling for age, sex, years of education and average food intake in gram per day. Subsequently, we reduced the number of observed ROIs according to our hypothesis, focusing on the ones located within the prefrontal and striatal areas (frontal pole (FP), superior/middle/inferior frontal gyrus (SFG/MFG/IFG), frontal orbital cortex (Forb), caudate nucleus, putamen, nucleus accumbens). This revealed significant negative correlations of daily tyrosine consumption and rsFC between bilateral putamen and left IFG pars opercularis [*β* = -0.35, *t*(280) = -3.30, *p-FDR* = .013], bilateral putamen and left SFG [*β* = - 0.33, *t*(280) = -3.07, *p-FDR* = .014], bilateral putamen and left IFG pars triangularis [*β* = - 0.30, *t*(280) = -2.92, *p-FDR* = .015], bilateral putamen and left MFG [*β* = -0.27, t(280) = - 2.44, *p-FDR* = .038], bilateral putamen and right IFG pars triangularis [*β* = -0.25, *t*(280) = - 2.43, *p-FDR* = .038] and bilateral putamen and right SFG [*β* = -0.24, *t*(280) = -2.34, *p-FDR* =.040](see *Table* 2 and *Figure* 2). We did not observe any significant tyrosine-mediated changes rsFC between caudate nucleus and prefrontal cortex, respectively between nucleus accumbens and prefrontal cortex. This suggests an association of higher tyrosine levels and a decrease in rsFC between striatal, in particular the bilateral putamen and prefrontal regions (SFG, MFG, IFG).

**Table 2:**
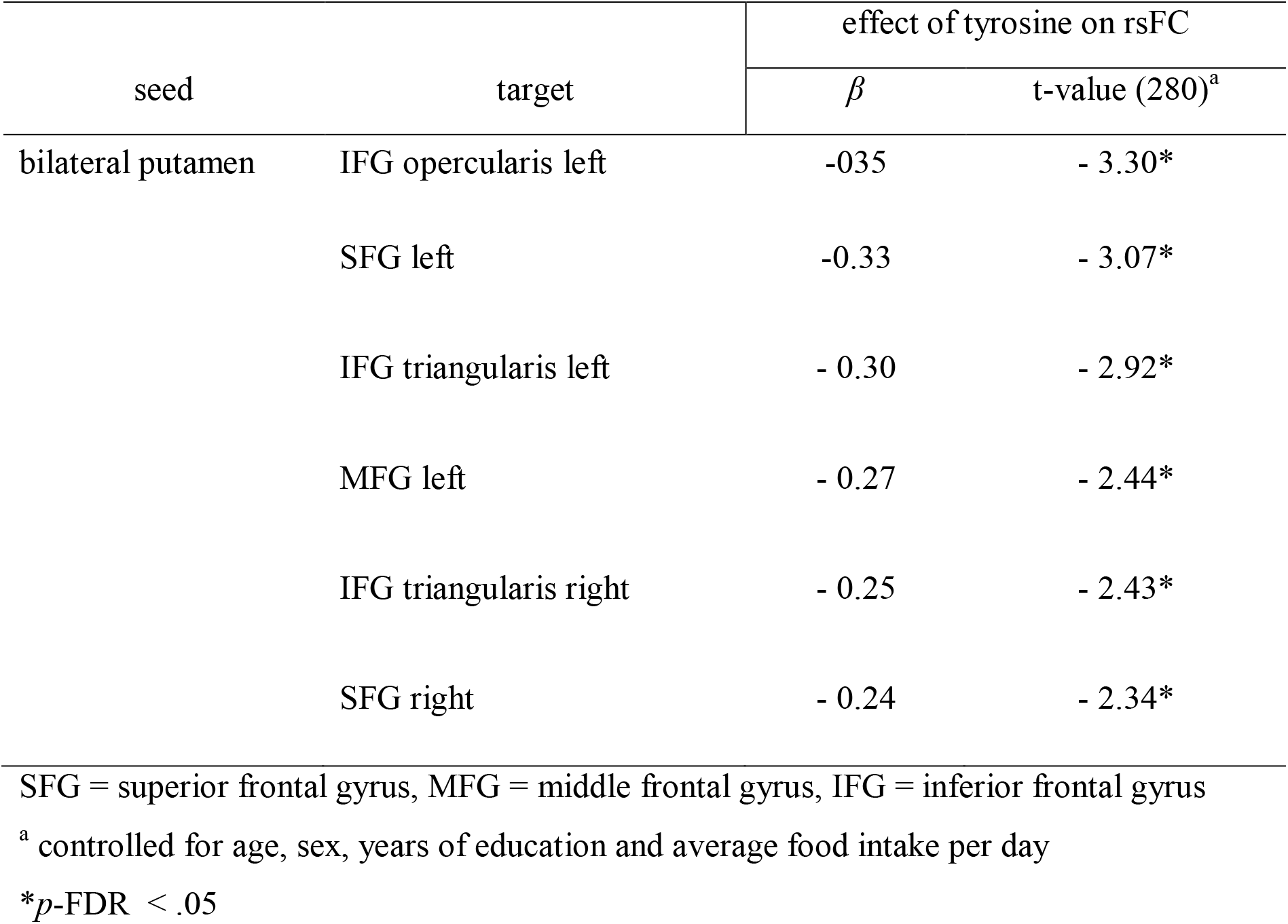
Effect of tyrosine intake per day on rsFC between striatal and prefrontal regions. Correlation of rsFC between striatal and prefrontal regions and WM performance.

**Figure 2:**
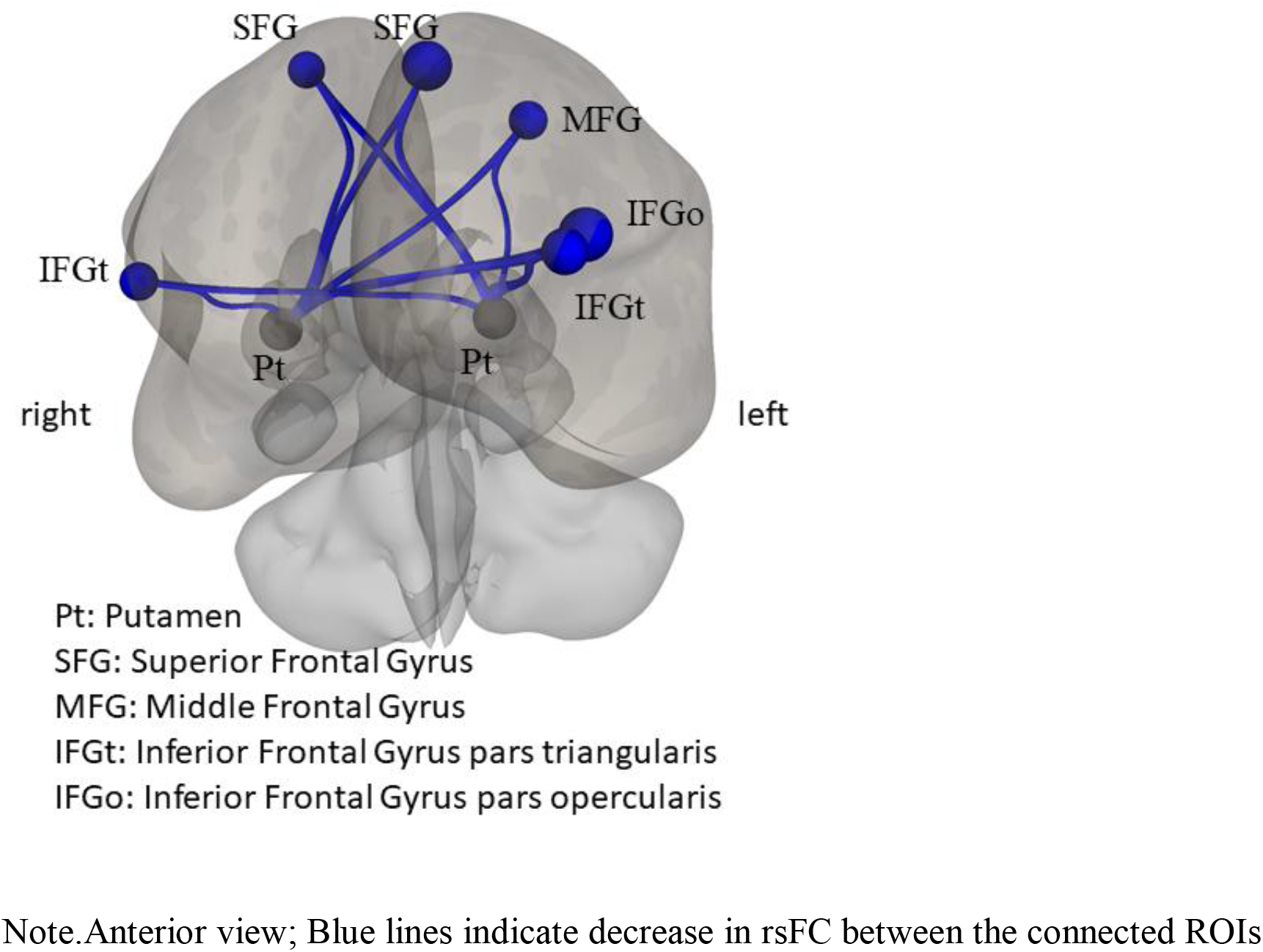
Correlations between daily tyrosine intake and rsFC between striatal and prefrontal regions

### Behavioral analysis

To analyze associations between daily tyrosine intake, WM and rsFC between brain regions revealed in the functional connectivity analysis areas, we first calculated the mean rsFC between these regions for each subject. The resulting parameter was the mean rsFC between the bilateral putamen and the bilateral SFG/MFG/IFG.

A multiple linear regression was calculated to predict WM based on average daily tyrosine intake in gram and mean rsFC. This regression model was significant [*F*(6, 279) = 9.096, *p*<.001], explaining 16,4% of the overall variance of WM. Average daily tyrosine intake significantly predicted WM (*β* = .209, *p* = .024), as did mean rsFC (*β* = -.134, *p* = .018). Subsequently, we performed a mediation analysis to investigate if rsFC mediates the effect of tyrosine on WM. Results indicated that tyrosine was a significant predictor of rsFC (*B* = -.257, SE = .079, *p* = .001), as well as rsFC a predictor of WM (*B* = -1.069, SE = .451, p =.019). Tyrosine remained a significant predictor of WM after controlling for the mediator (rsFC) (*B* = 1.373, SE = .605, *p* = .024), suggesting partial mediation. The indirect effect of tyrosine on WM mediated by rsFC was also significant (*B* = .274, SE = .129, CI = .080 –.625). These results suggest that some of the effects of average daily tyrosine intake on WM are mediated by differences in rsFC between the bilateral putamen and bilateral SFG/MFG/IFG. All analyses were controlled for age, sex, years of education and average daily food intake in gram. (see *Table* 3 and *Figure* 3)

**Table 3:** Effect of tyrosine on WM mediated by mean rsFC between bilateral putamen and the prefrontal cortex (bilateral SFG/MFG/IFG)

| Effect | *B* | SE | CI |
| --- | --- | --- | --- |
| tyrosine on mrsFC | -0.257 | 0.079* | - |
| mrsFC on WM | -1.069 | 0.451* | - |
| Direct effect:<br>tyrosine on WM | 1.373 | 0.605* | - |
| Indirect effect:<br>tyrosine on WM | 0.274 | 0.129<sup>a</sup> | 0.080 – 0.625 |

*B* = unstandardized coefficient, SE = standard error, CI = confidence interval, mrsFC = mean rsFC between bilateral putamen and bilateral SFG/MFG/IFG, WM = working memory Analysis was controlled for age, sex, years of education and average daily food intake in gram. Total effect [*B*(SE)] of tyrosine on WM = 1.647 (.599)\*.
\* $p < .05$
<sup>a</sup> $p$ is not calculated for the indirect effect

**Figure 3:**
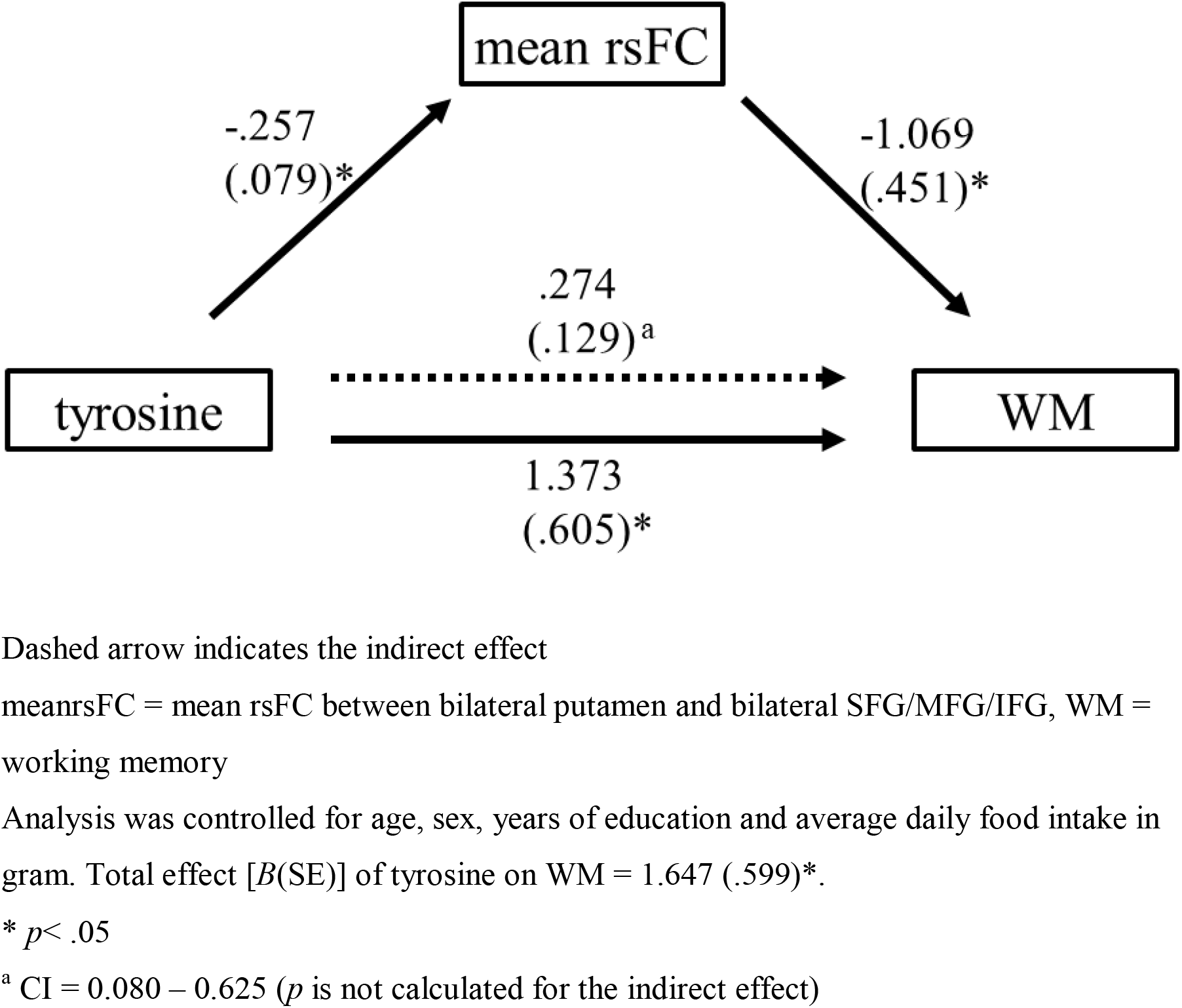
Effect of tyrosine on WM mediated by mean rsFC between bilateral putamen and the prefrontal cortex (bilateral SFG/MFG/IFG)

## Discussion

Recent research was able to show evidence for positive effects of dietary tyrosine intake on cognitive function in healthy humans (Kühn et al., 2017), which were similar to the ones observed after single high dose intake of tyrosine (Hase et al., 2015, Jongkees et al., 2015). Such results are explained by precursor availability – brain levels of tyrosine – influencing the synthesis rate of CA and therefore CA concentration within and around the neurons (Wurtman et al., 1974, Fernstrom et al., 2017). These precursor-mediated improvements in cognitive function were also accompanied by changes in functional connectivity (rsFC) between PFC and striatum (Nagano-Saito et al., 2008).

Based on the research mentioned above, we hypothesized an effect of nutritional tyrosine on functional brain connectivity, which could explain the observed improvements in WM capacity. We used data of the Berlin Aging Study II, in particular, resting-state fMRI scans and nutritional data derived from questionnaires to evaluate the hypothesized associations between diet and rsFC. Furthermore, results of WM dependent tasks were gathered to estimate a latent factor for WM. This latent factor capturing global WM abilities enabled us to testfor correlations between WM and daily tyrosine consumption, as well as associations with rsFC. We focused our research on WM, due to tyrosine-mediated effects on WM being well described within the scientific literature.

In the present study, we provided evidence for an association between dietary tyrosine intake and functional brain connectivity. These associations were also linked to positive effects on a latent factor for WM. Our findings are in line with previous research. Using a subsample of healthy elderly humans, from the same data set used by Kühn et al. (2017), we were able to replicate our previous findings of higher daily tyrosine intake being associated with higher WM capacity. In addition, we found a significantly decreased rsFC between putamen and prefrontal cortex, in particular SFG/MFG/IFG, in association with higher tyrosine consumption. This decrease in rsFC was likewise associated with higher WM performance. Furthermore, a mediation analysis revealed partial mediation of rsFC on associations between daily tyrosine intake and WM. These results suggest that some of the tyrosine-related differences in WM capacity in healthy elderly humans may possibly be a consequence of alterations in functional connectivity between striatal and frontal regions, in particular the putamen and SFG, MFG and IFG. In addition, there seem to be other direct effects of tyrosine on WM, which are not captured by alterations of rsFC between our observed brain regions.

What seems rather counterintuitive is the fact that increased tyrosine supply leads to decreased rsFC, which in turn is associated with better WM performance. We explain this result as follows: Our results indicate that higher supply of tyrosine is associated with decreased rsFC. In turn, lower supply is associated with an increase in rsFC. We speculate that this increase might represent a state of hyperconnectivity in subjects with insufficient tyrosine intake. Lower or insufficient precursor (tyrosine) supply might lead to reduced DA signaling and as a consequence to hyperconnectivity between cortical and subcortical regions. Similar findings can be made in pathological states of DA depletion such as patients suffering early stages of parkinson’s disease (PD). PD patients off medication show hyperconnectivity in resting-state fMRI between basal and cortical regions (Kwak et al., 2010). This increased connectivity was mitigated by therapeutic administration of L-DOPA. These findings fall in line with other reports on conditions of DA depletion, which have shown increased synchronous activity between basal and cortical areas (Costa et al., 2006; Gatev et al., 2006). According to these studies, increased synchronous activity between basal ganglia and its associated networks is a consequence of the pathological state of DA depletion. This effect of DA modulation was also shown in healthy adults. Pharmacological intervention using sulpiride, a DA antagonist leading to lower DA transmission, resulted in increased connectivity between basal ganglia and thalamic and cortical areas (Honey et al., 2008). Therefore, hyperconnectivity between such regions seems to represent a state of dysfunctional activity coordination, due to insufficient DA signaling. Higher age is commonly accompanied by DA depletion. Age-related decline of motor and cognitive functions is accompanied by impaired DA metabolism (Vernaleken et al., 2007; Erixon-Lindroth et al., 2005; Bäckman et al., 2000). At the same time, DA synthesis capacity seems to be upregulated in elderly humans (Braskie et al., 2008), possibly to compensate this reduction of signaling by an increase in DA synthesis. This upregulation of synthesis requires additional precursor supply, hence a higher demand for tyrosine can be expected. In conclusion, higher daily tyrosine supply seems to improve cognitive function within healthy elderly humans, due to mitigating increased synchronous resting-state activity between striatal and prefrontal regions. This dysfunctional activity coordination might be a consequence of age-related decline in DA metabolism. Higher tyrosine intake might be able to counteract this decrease in DA metabolism by increasing the synthesis rate. Our observations – in the context of former researchas mentioned in the introduction (Kühn et al., 2017)- suggest a greater need for tyrosine in healthy elderly humans to possibly improve cognitive aging by counteracting effects of age-related cognitive decline.

Dietary recommendations mostly focus on overall protein intake per day and have recently been updated to fit the higher demand of people older than 60 years. Research suggests a minimal daily intake of 1.0 to 1.2 g of protein (Bauer et al., 2013). According to this, we can expect a minimum average intake of roughly 80 g of protein per day. Within our study group 54.5% of participants consumed less than 80 g of protein, suggesting most people do not consume enough protein to satisfy their demand. Due to this, we think an increase in overall daily protein consumption, focusing on sources high in tyrosine like e.g. cheese (parmesan), soy, lean meats and fish, is an important way to address this problem, rather than taking supplements.

### Limitations

The present study is limited by the assessment time points of the data. Dietary data was gathered around two years prior to the cognitive data (mean difference in days = 719, *SD* =435) and MRI scans (mean difference in days = 824, SD = 430). Yet, we are convinced that the reports most likely reflect the participant’s regular diet, because the questionnaire focused on a rather long period of time, in particular the last 12 months. Moreover, we cannot exclude unobserved common causes for the association between tyrosine, differences in rsFC and cognition, which would explain our observations, due to the limitations of observational data. For example, there might be other nutrients frequently ingested within a tyrosine rich diet. However, due to the amount of research showing tyrosine-mediated improvement of cognitive function, we think tyrosine itself is most likely the key factor responsible for our observations. In addition, socio-economic status may play an important role explaining the observed associations. Higher socio-economic status is also associated with higher cognitive performance, as well as higher use of nutritional supplements, adherence to certain diet rules and lower fat intake (Hulshof et al., 1991). To control for this possible confound, we controlled for years of education as a rough approximation for the socio-economic status. However, the results were not affected by this variable. Measuring actual blood levels of tyrosine might also be interesting, to get a more accurate idea of the subject’s tyrosine supply. Therefore, we encourage future research to work in this direction, while also investigating the covariance structure of different nutrients. This may allow a deeper understanding of the role of tyrosine in cognition and high age. In addition, this might also allow us to give nutritional advice to improve cognition in healthy elderly humans.

## Informed consent

Informed consent was obtained from all individual participants included in the study.

## Conflict of interest

The authors declare no conflict of interest.

## Acknowledgements

We are grateful for the assistance of the MRI team at the MPI Berlin consisting of Sonali Beckmann, Nils Bodammer, Thomas Feg, Sebastian Schröder, Nadine Taube.

This article uses data from the Berlin Aging Study II (BASE-II) which has been supported by the German Federal Ministry of Education and Research under grant numbers 16SV5537/16SV5837/16SV5538/16SV5536K/01UW0808/01UW0706. The Responsibility for the contents of this publication lies with its authors.

SK has been funded by two grants from the German Science Foundation (DFG KU 3322/1-1, SFB 936/C7), the European Union (ERC-2016-StG-Self-Control-677804) and a Fellowship from the Jacobs Foundation (JRF 2016-2018). MB and CH have been funded by the German Science Foundation (SFB 936/C7).

